# Sex differences in cortisol and memory following acute social stress in amnestic mild cognitive impairment

**DOI:** 10.1101/757484

**Authors:** Kelly J. Murphy, Travis E. Hodges, Paul A.S. Sheppard, Angela K. Troyer, Elizabeth Hampson, Liisa A.M. Galea

## Abstract

**Objective:** Older adults with amnestic mild cognitive impairment (aMCI) develop Alzheimer’s type dementia approximately ten times faster annually than the normal population. Adrenal hormones are associated with aging and cognition. We investigated the relationship between acute stress, cortisol, and memory function in aMCI with an exploratory analysis of sex.

**Method:** Salivary cortisol was sampled diurnally and during two test sessions, one session with the Trier Social Stress Test (TSST), to explore differences in the relationship between cortisol and memory function in age-normal cognition (NA) and aMCI. Participants with aMCI (n=6 women, 9 men; mean age=75) or similarly aged NA (n=9 women, 7 men, mean age=75) were given tests of episodic, associative, and spatial working memory with a psychosocial stressor (TSST) in the second session.

**Results:** The aMCI group performed worse on the memory tests than NA as expected, and males with aMCI had elevated cortisol levels on test days. Immediate episodic memory was enhanced by social stress in NA but not in the aMCI group, indicating that stress-induced alterations in memory are different in individuals with aMCI. High cortisol was associated with impaired performance on episodic memory in aMCI males only. Cortisol in Session 1 moderated the relationship with spatial working memory, whereby higher cortisol was associated with worse performance in NA, but better spatial working memory in aMCI. In addition, effects of aMCI on perceived anxiety in response to stress exposure were moderated by stress-induced cortisol in a sex-specific manner.

**Conclusions:** We show effects of aMCI on Test Session cortisol levels and effects on perceived anxiety, and stress-induced impairments in memory in males with aMCI in our exploratory sample. Future studies should explore sex as a biological variable as our findings suggests that effects at the confluence of aMCI and stress can be obfuscated without sex as a consideration.

## Introduction

Normal aging results in declines in some cognitive domains, such as episodic memory, but not in others, such as experiential knowledge (Grady, 2012). Cognitive decline with aging is correlated with region-specific changes in prefrontal cortex and medial temporal lobe, a key change being hippocampal volume loss (Raz & Rodrigue, 2006) and these declines are accelerated in those with suspected Alzheimer’s Disease (AD) (Shi et al., 2009). Older adults with mild cognitive impairment (MCI) develop clinical dementia of the AD type at a rate of 10-30% annually, depending on MCI subtype, whereas those without MCI develop dementia at a rate of only 1% to 2% annually (Busse et al., 2003; Lupien et al., 1998). Thus, it is critical to identify neurobiological factors that may distinguish MCI from normal aging, such as differences in cortisol levels and their response to stress.

Dysregulation of the hypothalamic-pituitary-adrenal (HPA) system, including the stress hormone cortisol, has been linked to memory performance, aging, and hippocampal volume (Lupien et al., 1998; Sindi et al, 2014; Justice, 2018). Indeed, participants with suspected AD have higher levels of plasma (morning, 24 h release) or morning cerebrospinal fluid (CSF) cortisol than those experiencing normal cognitive aging (Hartmann et al., 1997; Doecke et al., 2012; Laske et al., 2011). Morning (CSF) levels of cortisol are also higher in MCI due to suspected AD, referred to as amnestic MCI (aMCI), as compared to older adults experiencing normal aging (NA) or MCI of other types (Popp et al., 2015). Moreover, aMCI individuals with higher morning CSF cortisol levels experienced accelerated clinical worsening and cognitive decline than those with a lower levels of cortisol, with a higher proportion of males in the aMCI group (Popp et al., 2015). Intriguingly, despite higher levels of cortisol, MCI participants experience lower levels of perceived stress during cognitive performance compared to age-matched healthy controls (Guerdoux-Ninot & Trouillet, 2019). Previous studies have rarely examined multiple timepoints of cortisol, stress induced cortisol, diurnal cortisol, or have used biological sex as a discovery variable, all factors which may contribute to the findings (Hidalgo et al., 2019; Yan et al., 2018). Cortisol is well known to vary in a diurnal pattern (Adam et al., 2017) and diurnal patterns are flattened in dementia of the AD type (Rasmuson et al., 2011; Ferrari et al., 2001). Importantly, to our knowledge, stress-induced cortisol has not been studied in relation to aMCI status and memory performance. Acute stress may be a more salient variable of HPA dysregulation to examine whether it can perturbate memory. Indeed, normally aging older adults who show less cortisol reactivity to acute stress using the Trier Social Stress Test (TSST) are more at risk to develop cognitive decline characteristic of MCI within 5 years (de Souza-Talarico et al., 2020). Thus, it is important to examine not only diurnal fluctuations in cortisol but also response to acute stress to determine whether these biomarkers modulate memory and how they may relate to progression to neurodegenerative disease. In addition, perceived stress, along with stress-induced cortisol, may be as important to investigate (Aggarwal et al., 2014). Furthermore, sex differences must be considered given previous studies have identified sex differences in cortisol levels in response to stress (Kudielka & Kirschbaum, 2005) which vary by age (for review see Hidalgo et al., 2019). Indeed, few studies to date have examined the relationships between cortisol, stress, memory, and potential differences between males and females in older populations (for review see Hidalgo et al., 2019), particularly with regards to MCI or AD.

Sex differences are seen in incidence of MCI (Gale et al., 2016; Koran et al., 2017; Mielke et al., 2014; Duarte-Guterman et al., 2020), with males more likely to develop MCI (both aMCI and non-amnestic subtypes) than females (Jack et al., 2019; Roberts et al., 2012), although there are conflicting reports that are likely due to methodological differences (e.g. Mielke et al., 2014). However, AD disproportionately affects females, with significant sex differences observed with regards to severity, neuropathological markers, and rates of cognitive decline (Irvine et al., 2012; Koran et al., 2017; Sohn et al., 2018). Sex differences in incidence of AD are not seen uniformly (Jack et al., 2019) and may depend on geographic location (Nebel et al., 2018) or greater longevity in women (Mielke et al., 2014). However, there are other sex differences in MCI to AD progression and symptom severity (Sundermann et al., 2017). For example, women tend to develop MCI at a later age, perhaps benefitting from their established superior verbal memory (Sundermann et al., 2017), but progress to AD more rapidly than men when adjusted for age (Irvine et al., 2012; Sohn et al., 2018). Sex differences in the trajectory of cognitive decline are also seen in MCI (Koran et al., 2017; Sohn et al., 2018). Indeed, females with more AD-associated neuropathology (total-tau and amyloid-beta [A□42] in cerebrospinal fluid), show greater declines in hippocampal volume and cognition compared to males, particularly among MCI individuals using the ADNI database (Koran et al., 2017; Sohn et al., 2018). Furthermore cognitive differences in verbal learning, delayed recall, visual learning and memory between NA and MCI females were significantly greater than those between NA and MCI males (Gale et al., 2016). This pattern in sex differences persisted in those with AD (Gale et al., 2016). Thus, sex differences in severity and progression to AD are seen in individuals with MCI and identifying the biological causes of this phenomenon is critical to treatment and prevention.

Gonadal production of sex steroids is reduced, but not entirely eliminated, with age in women and, to a lesser extent, men; however, adrenal cortisol production increases with age (Laughlin & Barrett-Connor, 2000). Although both sexes show increased cortisol levels with increased age, this effect is 3 times more pronounced in females (Otte et al., 2005) and increased cortisol levels are linked to poorer cognition and smaller hippocampal volume in older age (Lupien et al., 1998). Furthermore, as mentioned above females with MCI and AD present with greater declines in hippocampal atrophy and cognition than males (Irvine et al., 2012; Koran et al., 2017; Sohn et al., 2018), highlighting that the underlying pathophysiology of AD may be different in men and women and should be further explored. Although sex differences in AD have been identified, studies are scarce and even more so in MCI groups.

In this study we explored the relationship between diurnal fluctuations in cortisol, stress-induced cortisol, and memory performance in older adults experiencing aMCI and NA. A spatial working memory task known to be reliant on the integrity of the prefrontal cortex (Courtney et al., 1998) and an episodic and associative memory task known to be reliant on hippocampal integrity (Eichenbaum, 2017) were selected based on the number of glucocorticoid receptors and therefore sensitivity to cortisol in these brain regions (Dedovic et al., 2009) and the potential of fluctuations in cortisol to influence cognitive efficiencies. Because little is known as to whether there are sex differences in the relationship between aMCI and cortisol, we also used exploratory analyses of sex effects in the present study. We hypothesised that stress-induced cortisol, *via* the application of a psychosocial stressor (Trier Social Stress Test; Kirschbaum et al., 1993), would worsen memory scores and possibly alter perceived anxiety in individuals with aMCI compared to NA and that there would be sex differences in these effects.

## Materials and methods

### Participants

Older adults with age-normal memory (normal cognitive aging-NA) and with mild memory decline (aMCI) suggestive of neurodegenerative disease of the Alzheimer type (Albert et al., 2011) were recruited for this study and provided informed voluntary consent to participate. The following brief battery of neuropsychological measures were administered during Session 1 to confirm group membership: cognitive screening (Mini-Mental Status Exam (MMSE, Folstein & Folstein, 1974), expressive vocabulary (Vocabulary, Wechsler, 1997), attention tests involving auditory attention span (Digit Span, Wechsler, 1997) and speed and attention switching (Trail Making Tests A and B; -Spreen & Strauss, 1998), confrontation naming (Boston Naming Test, Kaplan et al., 1983), visuospatial construction, and immediate memory (Rey-Osterrieth Complex Figure-copy and immediate recall, Spreen & Strauss, 1998); and mood status (Hospital Anxiety and Depression Scale, HADS, Zigmond & Snaith, 1983). Criteria for establishing aMCI status were adherent to those described in Petersen (2004) and Albert et al. (2011). Participants were classified with single domain amnestic MCI if memory performance was revealed to be the only cognitive domain among those tested (which included attention, psychomotor speed, memory, language, visual spatial ability, and executive function) for which age-scaled scores were lower than expected based on estimated verbal IQ (established on a test of expressive vocabulary) and based on demonstrated performance in the other cognitive domains examined (see Table 1).

**Table 1.** Descriptive Data for the Participant Groups.

|  | NA<br>(*n* = 15) | aMCI<br>(*n* = 16) | Cohen's *d* |
| --- | --- | --- | --- |
| Age (years) | 75.3 (8.7) | 74.6 (8.0) | 0.08 |
| Female:Male ratio | 9:7 | 6:9 |  |
| Education (years) | 14.4 (3.5) | 14.9 (3.0) | 0.15 |
| MMSE | 29.0 (1.0) | 27.7 (1.9) | 0.89 |
| Vocabulary SS | 13.6 (2.9) | 13.8 (2.7) | 0.07 |
| Digit Span Forward SS | 11.9 (3.4) | 10.8 (2.9) | 0.34 |
| Digits Span Backward SS | 13.3 (3.3) | 12.7 (2.1) | 0.01 |
| TMT A SS | 10.3(1.48) | 9.47(1.92) | 0.48 |
| TMT B SS | 12.69(1.89) | 10.79(1.93) | 0.99 |
| Rey-Osterrieth Copy SS | 8.94(2.14) | 8.25(1.54) | 0.37 |
| Rey-Osterrieth<br>Immediate Memory* SS | 11.19(3.63) | 7.25 (2.63) | 1.24 |
| Boston Naming* SS | 12.68(2.75) | 10.80(3.53) | 0.59 |
| HADS Depression Scale | 2.6 (2.3) | 2.6 (2.3) | 0 |
| HADS Anxiety Scale | 5.1 (3.8) | 5.6 (2.5) | 0.16 |

*Note.* Mean scores with standard deviations in parentheses. NA= normal aging; aMCI = amnesic mild cognitive impairment; MMSE = Mini-Mental Status Exam; Vocabulary = Expressive vocabulary; Digits Span = Total attention span score for digits forward and backwards; TMT = Trail Making Test A (number sequencing) and B (alternating number-letter sequencing); Rey-O Copy = visual construction of the Rey-Osterreith Complex Figure; Rey-O Memory = immediate recall of the complex figure; SS = age-corrected scaled score. HADS = Hospital Anxiety Depression Scale with scores < 7 considered within normal limits. \*indicates significantly different between NA and aMCI

#### Normal Cognitive Aging (NA) group

Fifteen older adults (age 61-86 years) experiencing NA were recruited *via* community talks, newspaper advertisements, and databases of research volunteers. Prior to invitation to participate, normal general cognitive status, using the Telephone Interview for Cognitive Status, and health status were confirmed in a telephone screening interview. At the first of two sessions, health history was further queried to verify that there was no history of a neurological, medical, or psychiatric disorder, substance abuse, or medications affecting cognition. As described, NA was confirmed during Session 1 by measuring performance on a brief battery of neuropsychological tests. Three participants initially recruited as NA were identified as meeting criteria for aMCI based on Session 1 interview and lower than age and education expected performance on immediate recall of a complex figure.

#### Amnestic Mild Cognitive Impairment (aMCI) group

Sixteen individuals (age 59-85 years), recruited from physician referrals, from databases of research volunteers, and from newspaper advertisement, were classified as meeting the National Institute on Aging-Alzheimer’s Association classification criteria for aMCI (Albert et al., 2011). The single domain aMCI status of most of the aMCI participants had been previously established and the stability of this classification was confirmed by the interview and neuropsychological testing administered during Session 1. As previously stated, three of the participants initially recruited to the NA group were found to meet criteria for aMCI at Session 1 and were transferred to the aMCI group.

### Procedure

Participants completed alternate versions of episodic, associative, and spatial memory tasks across two test sessions (Figure 1) conducted 7-14 days apart. The TSST (Kirschbaum et al., 1993), a psychosocial stressor, was applied during the second test session and is described below. Salivary cortisol was collected during both test sessions for all participants and diurnal samples were collected as described below. Saliva was chosen because it most closely represents bioavailable cortisol (i.e., the fraction of the circulating hormone that is biologically available to tissues).

**Figure 1.**
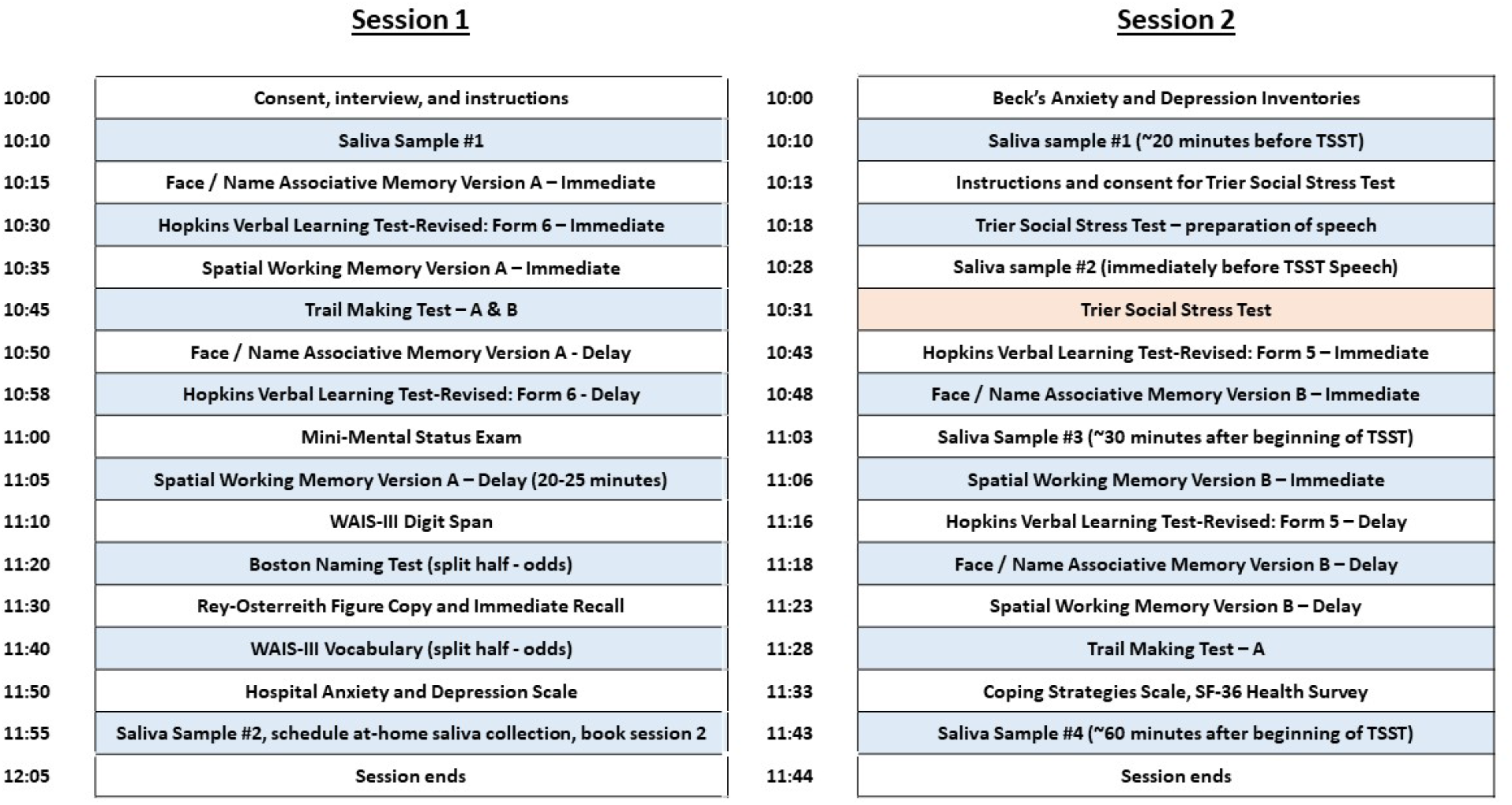
Timeline of Sessions and Testing within Sessions. Exact times varied based on individual variability.

#### Episodic memory

Two highly correlated versions (forms 5 and 6) of the Hopkins Verbal Learning Test-Revised (HVLT-R) were used (Session 1: form 6; Session 2: form 5; with the exception of one aMCI participant to whom they were presented in the opposite order). This task involves an oral presentation of a 12-item word list over three learning trials, followed by a 20-minute delayed recall trial and a forced choice yes/no recognition trial. The recognition trial consists of 24 items comprised of the 12 target words, six semantically related foils, and six un-related foils. Measures of interest included total recall across three learning trials, total delayed recall, and retention.

#### Associative recognition

A face-name associative recognition test, created by Troyer and colleagues (2011, 2012; modeled after Mayes et al., 2004), was used. Stimuli consisted of visually presented black-and-white images of faces (half male and half female) paired with aurally presented first names. Two versions of the task were used, each with 28 face-name pairs. During the task, 20 faces were individually presented on the computer screen for 6 seconds each with an inter-stimulus interval of 0.5 seconds; the examiner read the name associated with each face at the onset of each new face stimulus. Two study phases, differing only in stimulus presentation order, were administered in succession because our previous research indicated that item memory and association memory differences increase after repeated learning trials (Troyer et al., 2008). Only 16 of the 20 face-name pairs were considered test items as the first and last pairs in each study phase presentation were excluded to reduce primacy and recency effects on recognition accuracy. Following a 30 second delay, yes/no recognition testing was conducted with 24 face-name pairs presented, including eight intact pairs, eight recombined pairs, and eight new pairs presented in random order. During testing the examiner orally presented the name in the form of a question “Did I tell you this was [NAME]?” when the face appeared on the screen. Participants were instructed to say “yes” only to faces they had seen before that were paired with the correct name and “no” to faces they had not seen before, faces that were paired with the wrong name, or names they had not heard before. Immediately following testing, procedure verification was undertaken (i.e. participants retold the yes/no rules to the examiner). Participants were presented with unique, but equivalent (Troyer et al., 2011), sets of face-name pairs during Sessions 1 and 2. Associative recognition calculated as the difference between the proportion of correctly identified intact pairs (same face-name pairs viewed at study) and proportion of false alarms to recombined pairs (different pairings of previously viewed faces and names at study) was the primary measure of interest. The decision to focus on associative recognition was based on our previous research demonstrating this measure was most sensitive to aMCI and hippocampal volume loss (Troyer et al., 2012).

#### Spatial working memory

This task was modeled after the stimuli and procedures of Duff and Hampson (2001). A 4×5 rectangular array, measuring approximately 27cm in length and 34cm in width, consisting of coloured squares (10 colours, each represented twice) that were hidden under removable covers, was presented on a tabletop at which participants were seated. The coloured squares were randomly arranged on a uniform white backing and completely concealed beneath uniform white covers that could be temporarily lifted by participants to reveal the coloured square beneath. Participants were instructed to find all 10 pairs of matching coloured squares in as few choices as possible by lifting the covers two at a time. Prior to beginning the task, participants were familiarised with the colours of the test stimuli by having them view and name a set of 10 individual coloured squares. Each time a matching pair was located on the stimulus array, the examiner placed an individual coloured square representing the colour of the pair discovered at the top of the rectangular array, so participants did not need to remember which colour pairs had been found. Measures of interest included: the number of choices (squares uncovered) made in discovering all 10 matching pairs (criterion) and the time taken to reach criterion. Participants were told they would be timed and that they should attempt to locate all 10 pairs in as few choices as possible. Once they reached criterion, a second trial was immediately administered, with a third trial administered following a 30-minute delay. Locations of the coloured squares was constant within session but changed between Sessions 1 and 2.

#### Psychosocial stressor (TSST)

The psychosocial stressor used in this study was modeled after the TSST (Kirschbaum et al., 1993). The stressor was introduced immediately following the first saliva collection at 10:10h. Participants were instructed to prepare a five-minute speech on the topic of ‘ *The effect of tuna fishing on the dolphins and other ocean animals’* to be presented to a panel of three evaluators, including the examiner. They were given a pencil and paper and told to write down the points they would like to make in their speech for which they would have 10-minutes to prepare. The examiner then left the room and returned 10 minutes later, collected a saliva sample from the participant (anticipation period), and then led the participant to a conference room to give their speech. Participants were instructed to leave their written notes behind, to give their speech from memory, and to try and speak for five minutes, which was timed by the examiner. Immediately following the public speech, participants engaged in a five-minute serial subtraction task in which they were asked to count backwards aloud by 13 from the number 1022 as quickly and accurately as possible in front of the panel of evaluators. When an error was committed the participant was instructed to begin again from the number 1022. Because perceived stress/anxiety has been found to be lower in aMCI compared to NA individuals performing an attention-based task (Stroop-task; Guerdoux-Ninot & Trouillet, 2019), participants were asked if they found the test anxiety provoking and if so to rate their perceived anxiety during the public speech from 0 (low anxiety) to 100 (high anxiety). Lastly, participants were led back to the test room to undertake memory testing (episodic, associative, spatial) and to provide additional saliva samples as described below.

### Saliva collection and analysis

Multiple salivary cortisol samples were taken using the Salivette™ method (Sarstedt Inc, Sarstedtstraße, Numbrecht, Germany) across both the test sessions (for all participants) and at home (to examine diurnal variations). Participants were instructed not to eat, drink, or smoke for at least 30 minutes prior to saliva collection and to rinse their mouths with water 5 minutes before collecting the saliva. For the test sessions, both sessions were conducted in the morning from 10:00h to 12:00h. Saliva was collected at 10:10h and 12:00h at the first test session and during the second session at 10:10h; immediately following the anticipation period to the application of a psychosocial stressor (~10:30h); 30 minutes following the application of the psychosocial stressor (~11:00h); and at about 12:00h at the conclusion of the second test session. In addition, basal salivary diurnal cortisol samples were requested from participants across three agreed upon days intervening between the test sessions. Five samples were collected per day on the following schedule: 30-minutes after awakening (ranged from 5:30 am to 8:30h), 09:00, 16:00, 19:00, and 21:00h. Phone call reminders and verifications were provided by the examiner for each of the 4 specified clock times on all three collection days. Participants were given pre-labeled saliva collection tubes at the end of Session 1 and instructed to store their collected samples in the home refrigerator. Collection time of day was further verified by requiring participants to record the collection time on a label provided on the collection tube. The basal samples were gathered from participants when they returned for the second test session.

Saliva was centrifuged at 1500g and kept frozen at −20°C prior to analysis. Cotton-based collection is suitable for cortisol determinations (Büttler et al., 2018). Salivary cortisol was analyzed in duplicate by the Neuroendocrinology Assay Laboratory at the University of Western Ontario (EH). An established ^125^I solid-phase radioimmunoassay was used (Norman et al., 2010), based on antibody and tracer obtained from Siemans Healthcare Diagnostics (Deerfield, IL). The Laboratory specialises in saliva determinations. Briefly, saliva was analyzed directly, without extraction, using a 200μL aliquot and an extended 3hr incubation at room temperature. The calibration curve was diluted 1:10 and ranged from 0-138 nmol/L. The intra-assay coefficient of variation calculated across low, medium, and high pools averaged 4.2% and the sensitivity of the assay was < 0.25 nmol across 3 assay runs. All samples from a given participant were analyzed in the same assay run and the average salivary cortisol concentration across the two duplicates (in nmol/L) was the value used for all statistical analyses. All cortisol data was log-transformed (log10) due to non-normality.

We also examined the diurnal and TSST cortisol levels area under the curve (AUC) using two formulas for AUC one with respect to the ground (AUCg) and one with respect to the increase (AUCi) (Pruessner et al., 2003) using the following formulas:

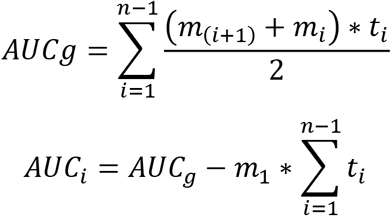

Where t_i_ – is the distance between time measurements), m_1_ – is the first cortisol measurement

### Data Analyses

Statistical analyses were performed using Statistica TIBCO Software (Palo Alto, CA, USA) and SPSS with the PROCESS package. Because we were examining sex differences as an exploratory factor, we initially conducted analyses on age and education level between the sexes (see Table 2). Due to differences in ages between groups we used age was used as a covariate in all analyses. General linear model repeated measures analyses of covariance (RMANCOVAs) were conducted on log-transformed cortisol measures (diurnal, session time) and cognitive measures (HVLT, SPWM, AR) using group (NA, aMCI) and sex (male, female) as the between-subject factors and with either Session (1, 2) or time of day (30 min, 9 AM, 4PM, 7pm, 9PM) or time during test Session (10:10, 10:30, 11, 1130) as the within-subjects factor. On certain measures (represented in Table 1) general linear model ANCOVA were conducted using group (NA, aMCI) as the between-subject factor. *Post hoc* analyses used Newman-Keul’s comparisons. Due to the small sample size, exploratory analyses on possible sex effects were run using a Bonferroni correction on *a priori* analyses, using a two-tailed significance criterion of p=0.10. Otherwise statistical significance was p=0.05 was used for all other statistical tests conducted. Chi-square analyses were completed for frequency of sexes between groups and for anxiety scores. For exploratory moderation analyses hierarchical linear regressions were performed to test the overarching hypothesis that the magnitude of Session 1 and Session 2 cortisol moderates the relationship between group (MCI, NA) and perceived anxiety during the TSST, episodic memory, associative memory, or working memory. Hierarchical linear regressions that included either Session 1 cortisol (AUCg) or Session 2 cortisol (AUCg) as a moderators were conducted for males and females separately. Dummy codes were created for participant group (MCI = 1, NA = 0), cortisol data were log10 transformed and standardized, and all other data were standardized. Regression models with significant interactions are reported.

**Table 2:** Demographic Data for the Participant Groups.

| | NA<br>males<br>(*n* = 7) | aMCI<br>males<br>(*n* = 9) | NA<br>females<br>(*n* = 9) | aMCI<br>females<br>(*n* = 6) | P values | Effect Size:<br>$\eta^2_p$ |
| --- | --- | --- | --- | --- | --- | --- |
| Age<br>(years)* | 71.3<br>(8.8) | 76.0 (7.7) | 79.8<br>(5.9) | 71.5<br>(8.0) | Sex ns<br>Group ns<br>Group by sex<br>0.025 | Sex .02<br>Group .02<br>Group by sex .017 |
| Education<br>(years) | 14.7<br>(3.3) | 15.5 (2.2) | 14.1<br>(3.6) | 13.5<br>(2.7) | Sex ns<br>Group ns<br>Group by sex<br>*ns* | Sex .05<br>Group .00<br>Group by sex .01 |

*Note.* Mean scores with standard deviations in parentheses. NA= normal aging; aMCI = amnesic mild cognitive impairment \* denotes significant difference.

## Results

### Males with aMCI had higher levels of cortisol than NA males during the test sessions

Cortisol levels during the two test sessions were analyzed separately as the psychosocial stressor (TSST) was conducted during Session 2. For Session 1 (without the TSST), males with aMCI had significantly higher levels of cortisol than any other group at the first time point (10:10AM) (all *p*s<0.001; Cohens d: for NA males=0.79, NA females=1.99, and aMCI females=1.67); time by group by sex interaction: F_(1, 27)_=3.65, *p*=0.066, η_p_=0.12; Figure 2A). There were also main effects of sex and time (main effect of sex: F_(1,27)_=16.24, *p*<0.001, η_p_=0.37; main effect of time: F(_(1,27)_=19.515, *p*<0.0001, η_p_=0.42). For AUCg in Session 1, aMCI males had higher levels of cortisol than all other groups (all *p*s<0.05; Cohens d: for NA males=0.81, NA females=1.57, and aMCI females=1.95; interaction effect of sex by group: F_(1, 27)_=4.013, *p*=0.055, η_p_=0.129; Figure 2B). There was also a main effect of sex (F_(1,27)_=10.25, *p*=0.003, η_p_=0.28) but not group (*p*=0.37; see Figure 2B). There were no significant differences using AUCi (*p*>0.15, group: η_p_=0.03, sex η_p_=0.06, group by sex η_p_=0.07).

**Figure 2.**
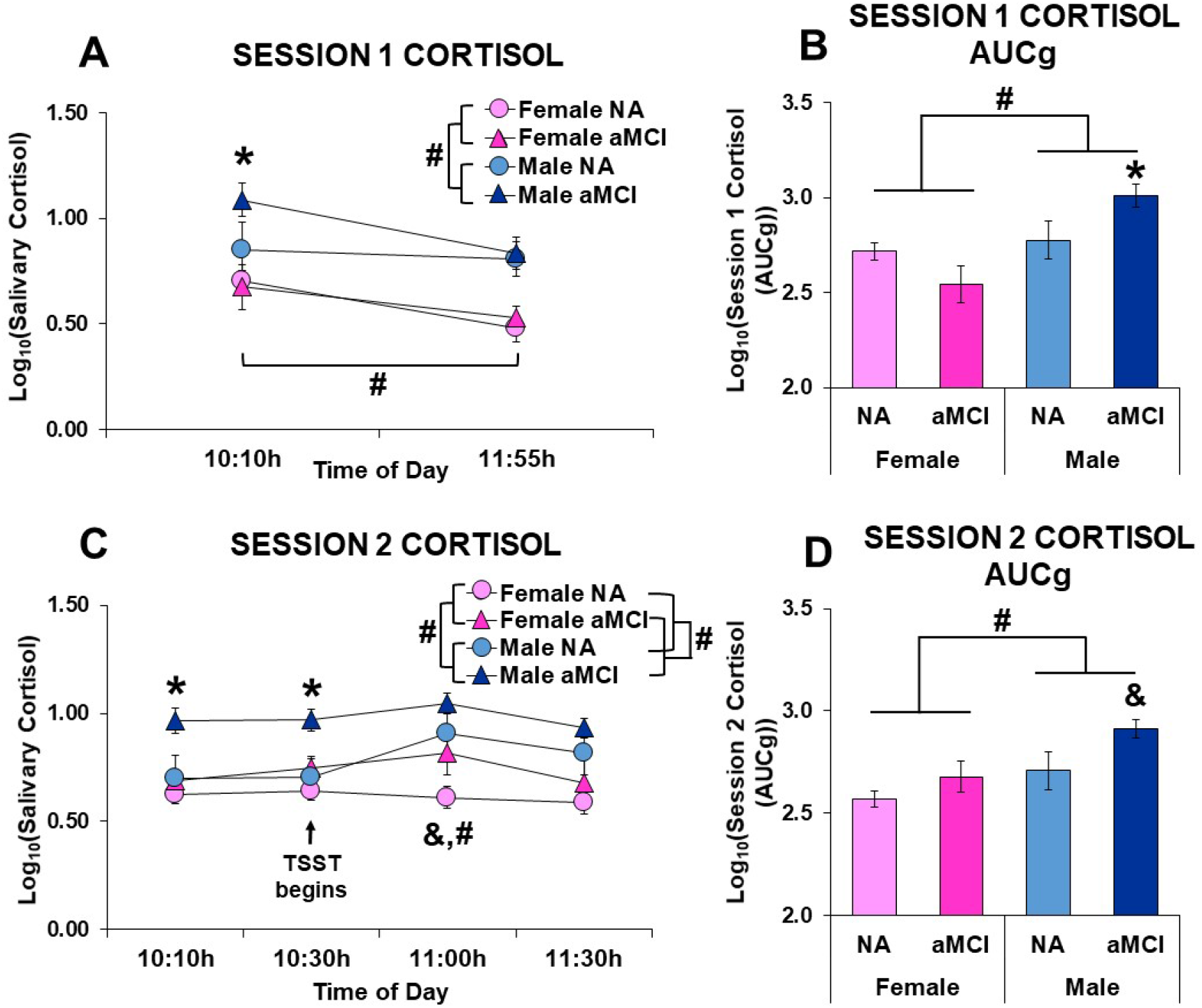
(A) Log_10_ transformed salivary cortisol at two separate time points and (B) Log10 transformed area under the curve with respect to the ground (AUCg) cortisol during Session 1 in normal aging (NA; *n* = 9 females, *n* = 7 males) and amnestic mild cognitive impaired (aMCI; *n* = 6 females, *n* = 9 males) participants. (C) Log transformed salivary cortisol at four separate time points and (D) Log10 transformed AUCg cortisol during Session 2 in NA (*n* = 9 females, *n* = 7 males) and aMCI (*n* = 6 females, *n* = 9 males). # denotes a Main effect of sex, group, or time, *p* < 0.05. * denotes aMCI males differ from all other groups at a single time point or compared to all other groups for AUCg cortisol, *p* < 0.05. & denotes aMCI females or males differ from NA females or males respectively at a single time point, *p* < 0.05.

During Session 2, in which the TSST was performed, aMCI participants had higher cortisol levels than NA (main effect of group: F_(1,27)_=7.86, *p*<0.009, η_p_=0.23). Furthermore, males had higher levels of cortisol than females (main effect of sex: F_(1, 27)_=13.58, *p*=0.001, η_p_=0.33). Furthermore, cortisol levels were highest 30min after the TSST was initiated compared to all other time points as expected (all *p*s<0.03; main effect of time: F_(3,81)_=3.95, *p*=0.01, η_p_=0.13). *A priori* analyses indicated that males with aMCI had significantly higher levels of cortisol during the first two time points in the second session prior to the TSST than all other groups (all *p*s <0.001; Cohens d: timepoint 1: for NA males=1.45, NA females=2.33, and aMCI females=2.00, timepoint 2: for NA males=1.25, NA females=2.44, and aMCI females=1.82). Importantly, males with aMCI did not show any changes in cortisol levels across Session 2 (all *p*s >0.18) whereas females with aMCI had significantly higher levels of cortisol 30min post-TSST than NA females (*p*=0.003 Cohens d=1.03) but not at any other time point (all *p*s>0.18; Cohens d= 0.4-0.98; Figure 2C).

For AUCg, in Session 2 we found that males had higher levels of cortisol than females (main effect of sex: F_(1,27)_=8.36, *p*<0.007, η_p_=0.24) and aMCI had higher levels than NA (main effect of group: F_(1, 27)_=4.202, *p*=0.05, η_p_=0.135). A priori we also found that aMCI males had higher AUCg than NA (*p*=0.060, one-tailed, 0.03; Cohens d: for NA males=0.77, NA females=2.39, and aMCI females=1.40) which was not evident in the females (*p*=0.35; Cohens d=0.73 between female groups) see Figure 2D). There were no significant differences using AUCi (*p*>0.27, group: η_p_=0.00, sex η_p_=0.004, group by sex η_p_=0.044).

### TSST was endorsed as anxiety provoking by females more than males with aMCI

Both sexes in the NA group endorsed the TSST as anxiety provoking, with 57% of participants indicating that the TSST was anxiety provoking. However, among aMCI participants 80% of females but only 11% of males indicated the TSST speech was anxiety provoking (χ^2^=6.644, *p*<0.01, Table 2). Of those participants who rated the TSST as anxiety provoking there was no significant difference in the rating of the anxiety level (*p*s>0.43, main effect of group η_p_=0.001, main effect of sex= η_p_=0.002, interaction η_p_=0.07; ratings (mean and standard deviation) and sample size of those that found the TSST anxiety provoking: NA females (n=4)=68.25±25.8, aMCI females (n=4)=50.25±26.10, NA males (n=4)=49.75±35.9; MCI males(n=1)= 64).

### Social Stress enhanced immediate recall in the NA but not in participants with aMCI. aMCI participants performed worse on episodic, associative and spatial working memory than NA

Immediate recall of HVLT-R was enhanced after the TSST in Session 2 compared to Session 1 in the NA (*p*=0.004), but no such enhancement was seen in participants with aMCI (*p*=0.245; interaction: group by session: F_(1,27)_=7.6, *p*=0.010, η_p_=0.22, main effect of group: F_(1,27)_=41.4, *p*<0.001, η_p_=0.61). Breaking this down by sex, the NA groups, regardless of sex, showed enhanced recall in Session 2 compared to Session 1 (males (*p*=0.02, Cohens d= 1.14) females (*p*=0.04, Cohens d=0.61)). However, aMCI males had impaired immediate recall (*p*=0.04, Cohen’s d= 0.63) on Session 2 following the stressor compared to Session 1, with no significant enhancement in the females with aMCI across sessions (*p*=0.47; Cohen’s d=0.25, Figure 3A). This enhancement in recall following the TSST was also seen for delayed recall for the HVLT-R which was administered 40 minutes after the TSST with all groups scoring better on Session 2 than Session 1 (main effect of session F_(1,27)_=7.15, *p*=0.012, η_p_=0.21) and with aMCI scoring worse than NA (F_(1,27)_=63.22, *p*<0.001, η_p_=0.70; Figure 3B). For HVLT-R retention, aMCI participants performed worse compared to NA participants regardless of session (main effect of group F_(1,26)_=27.09, *p*<0.001, η_p_=0.51; Figure 3C).

**Figure 3.**
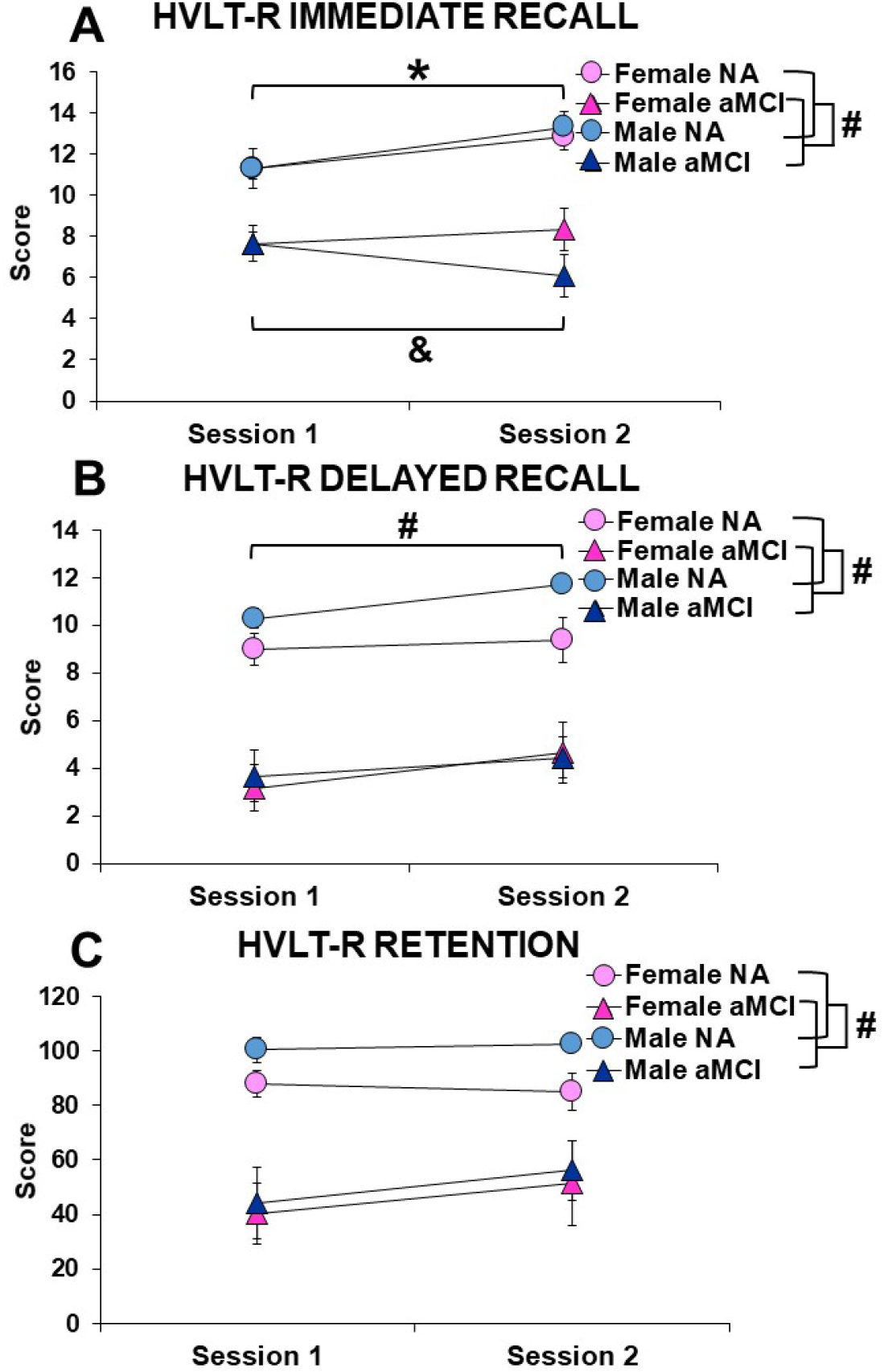
Hopkins Verbal Learning Test-Revised (HVLT-R) (A) immediate recall, (B) delayed recall, and (C) retention scores during session 1 and session 2 (after Trier Social Stress Test) in normal aging (NA; *n* = 9 females, *n* = 7 males) and amnestic mild cognitive impairment (aMCI; *n* = 6 females, *n* =9 males). # denotes Main effect of group or session, *p* < 0.05. * denotes a significant increase from Session 1 and Session 2 in NA males and females, *p* < 0.05. & denotes a significant decrease from Session 1 and Session 2 in aMCI males, *p* < 0.05.

For associative memory as expected aMCI participants remembered fewer face-name pairs than NA (main effect of group: F_(1,27)_=32.86, *p*<0.000, η_p_=0.55). There were no other main or interaction effects (all *p*s> 0.25).

During the spatial working memory across sessions and trials, aMCI participants made more choices than NA across all trials (main effect of group: F_(1, 27)_=13.6, *p*=0.001, η_p_=0.34). Furthermore, all participants required fewer choices by the delay trial (all *p*s<0.02; main effect of trial (F_(2,52)_=4.80, *p*=0.012, η_p_=0.16). There were no other significant main effects or interactions (all *p*s>0.20; all η_p_ <0.09; Figure 4B).

**Figure 4.**
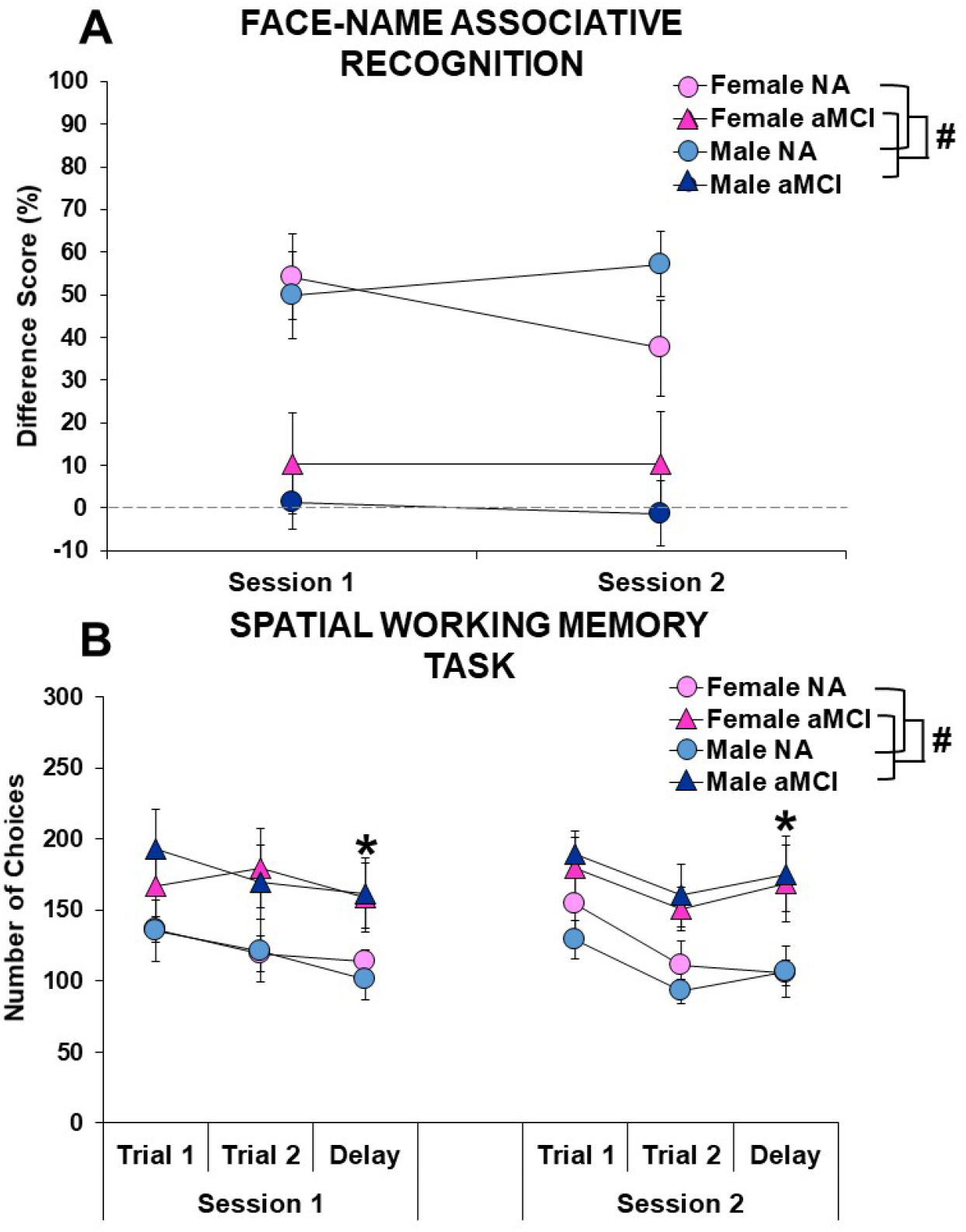
(A) Face-name associative recognition and (B) spatial working memory performance during Session 1 and Session 2 in normal aging (NA; *n* = 9 females, *n* = 7 males) and amnestic mild cognitive impaired (aMCI; *n* = 6 females, *n* = 9 males) participants. # denotes main effect of sex or group *p* < 0.05. * denotes fewer number of choices in delay trial compared to Trial 1 and Trial 2.

### Diurnal CORT did not differ between male groups

Only 21 of 31 participants completed all five time points for cortisol collection across three consecutive days. We used multiple imputation to calculate missing values for all individuals with more than 75% of the data available (2 participants only had 2/15 and one had 9/15 samples available). Curiously these 3 participants were females with aMCI. As we did not want to use the imputed values for so many missing datapoints we performed the analyses only with males. Analysis of these 16 participants revealed only a main effect of time (F_(4,56)_=48.51, *p*<0.001, η_p_=0.776; Figure 5A), but no main effect of group (F_(1,14)_=0.0349, *p*=0.85, η_p_=0.002 or interaction (F_(4,56)_=0.49, *p*=0.74, η_p_=0.033). *Post hoc* analyses revealed that cortisol was higher at each early timepoint than all other timepoints except there was no difference between the two evening samples. There were no significant differences between groups in time of awakening.

**Figure 5.**
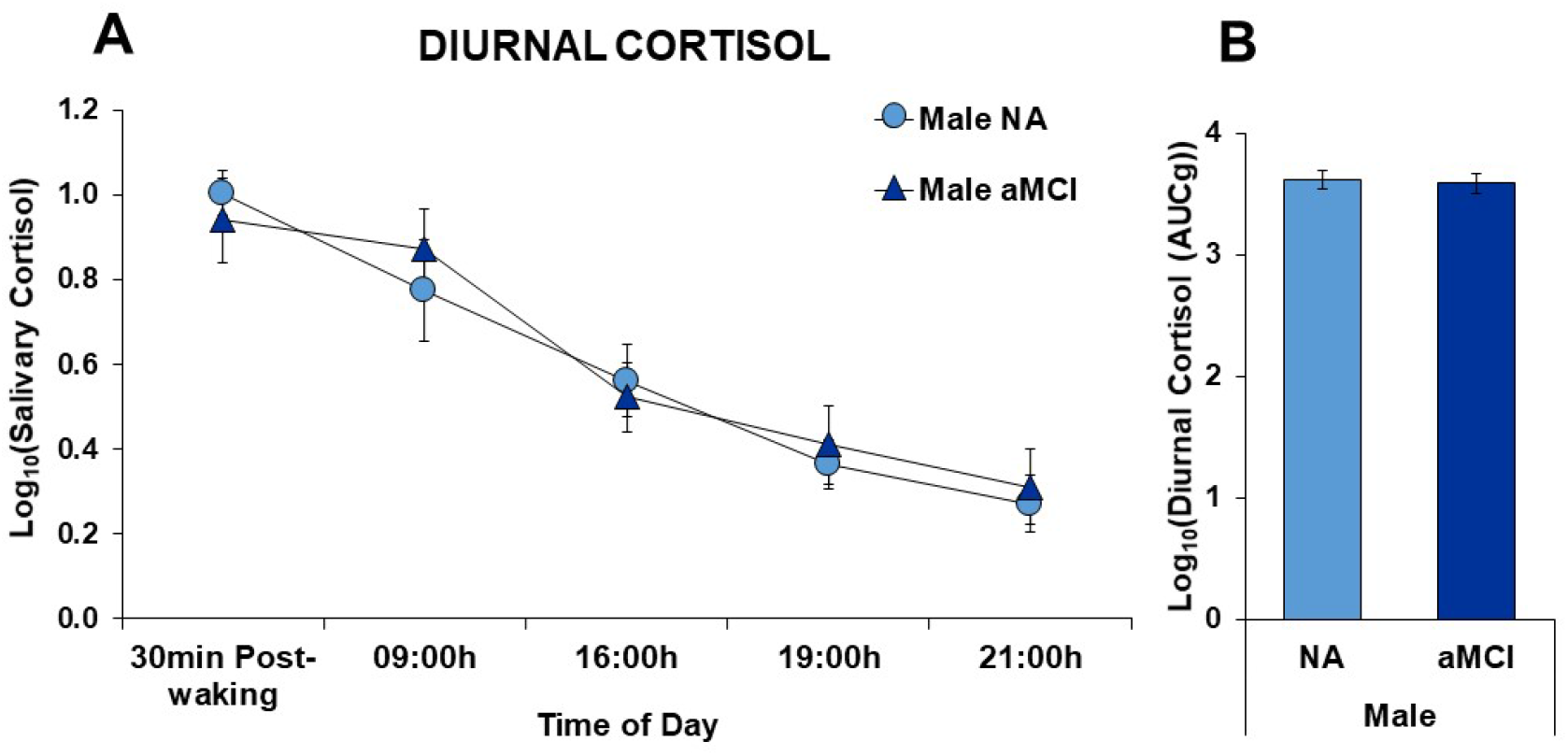
(A) Log_10_ transformed diurnal salivary cortisol measures at five time points averaged across three consecutive days in normal aging (NA) males (*n* = 7) and amnestic mild cognitive impairment (aMCI; *n* = 9). (B) Log transformed area under the curve with respect to the ground (AUCg) diurnal cortisol in NA and aMCI males.

We also calculated average cortisol AUCg and AUCi across all three days in males only. There were no significant effects for AUCg (*p*=0.549, η_p_=0.03) or AUCi (*p*= 0.6, η_p_=0.02).

### Session 1 Cortisol (AUCg) was associated with better spatial working memory in aMCI females, but the reverse in NA females

We next correlated Session 1 and Session 2 AUCg with episodic, associative and spatial working memory measures across group and sex, as we saw no significant differences in AUCi. Session 1 had the only significant correlations, with females showing positive associations with AUCg and better performance in spatial working memory in aMCI females (in trial 2 (r=-0.839, *p*=0.037) and the delay trial (r=-0.952, *p*=0.003)) but the opposite patterns in NA females (trial 2 (r=0.796, *p*=0.01)). These correlations were significantly different (z=3.261, *p*=0.001). In NA females, AUCg was positively associated with better HVLT-retention (r=0.7017, *p*=0.035), but negatively associated with associative memory (r=-0.703, *p*=0.035). In males, the only correlation was positive in Session 1 was associative memory and AUCg cortisol in aMCI males (r=0.818, *p*=0.007). There were no other significant correlations in males or for data in Session 2.

### Session 1 cortisol (AUCg) moderates the relationship between aMCI and episodic memory retention in females, working memory in both females and males, and associative recognition in males

We tested the hypothesis that the magnitude of cortisol (AUCg) moderates the relationship between participant group (aMCI, NA) and episodic (HVLT), associative (face-name), or working memory (spatial working memory) during Session 1 in males and females. Regressions revealed that higher cortisol predicted lower HVLT-R retention scores in aMCI females and higher HVLT-R retention scores in NA females (model: *F*_(3,11)_=11.64, *p*=0.001, *R^2^*=0.760, adjusted *R^2^*=0.695; interaction: *b*=-1.944, β=-1.124, t_(14)_=-5.653, *p*<0.001, *sr*^2^=0.696). Further, high Session 1 cortisol predicted a lower number of choices in aMCI females and a higher number of choices in NA females during trial 1 (model: *F*_(3,11)_=10.2, *p*=0.002, *R^2^*=0.735, adjusted *R^2^*=0.663; interaction: *b*=-1.527, β=- 1.562, t_(14)_=-4.413, *p*=0.001, *sr*^2^=0.468) and the delay trial (model: *F*_(3,11)_=11.21, *p*=0.001, *R^2^* 0.754, adjusted *R^2^*=0.686; interaction: *b*=-1.354, β=-1.228, t_(14)_=-3.593, *p*=0.004, *sr*^2^=0.289) of the spatial working memory task. In males, high Session 1 cortisol predicted increased associative recognition in aMCI males versus a decline in NA males (model: *F*_(3,12)_=9.723, *p*=0.002, *R^2^*=0.709, adjusted *R^2^*=0.636; interaction: *¿*=0.915, β=0.658, t_(15)_=2.466, *p*=0.030, *sr*^2^=0.148). High Session 1 cortisol also marginally predicted a lower number of choices in aMCI males versus NA males for spatial working memory during trial 2 (model: *F*_(3,12)_=2.320, *p*=0.127, *R^2^*=0.367, adjusted *R^2^*=0.209; interaction: *b*=-1.389, β=-0.856, t_(15)_=-2.177, *p*=0.050, *sr*^2^=0.250) (see Figure 6). Additionally, although no interaction was significant, group was a significant predictor for session 1 HVLT-R immediate recall and delayed recall in females and males and a predictor for face-name associative recognition in females (all *p*s<0.05) (see Table 3).

**Figure 6.**
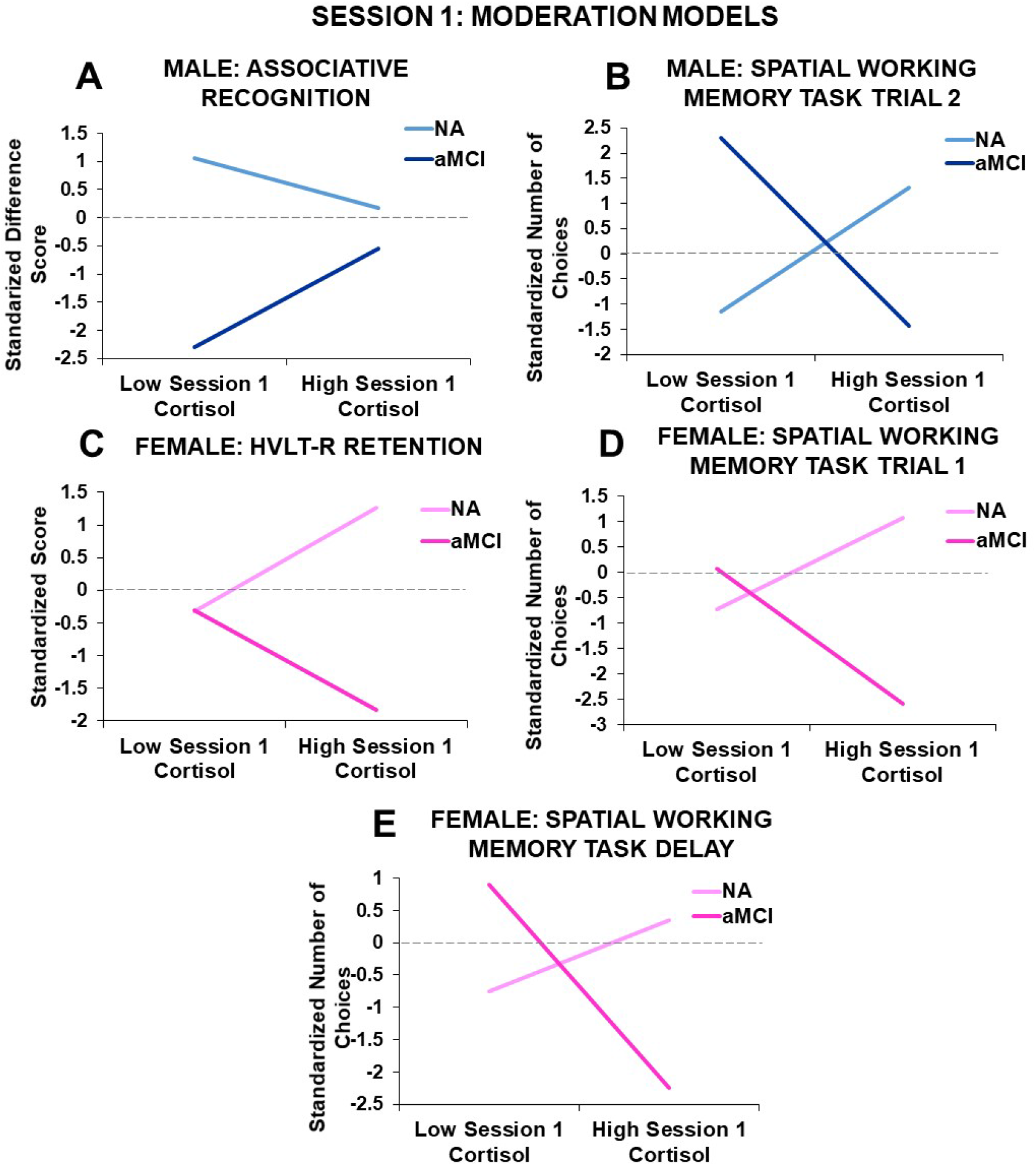
Standardized moderation effects of Session 1 cortisol (AUCg) on the relationship between participant group (normal aging (NA), amnestic mild cognitive impairment (aMCI)) and Session 1 (A) associative recognition scores and (B) spatial working memory task Trial 2 choices in males (*n* = 7 NA, *n* = 9 aMCI) and (C) Hopkins Verbal Learning Test-Revised (HVLT-R) retention scores, and (D) spatial working memory task Trial 1 and (E) delay choices in females (*n* = 9 NA, *n* = 6 aMCI).

**Table 3.**
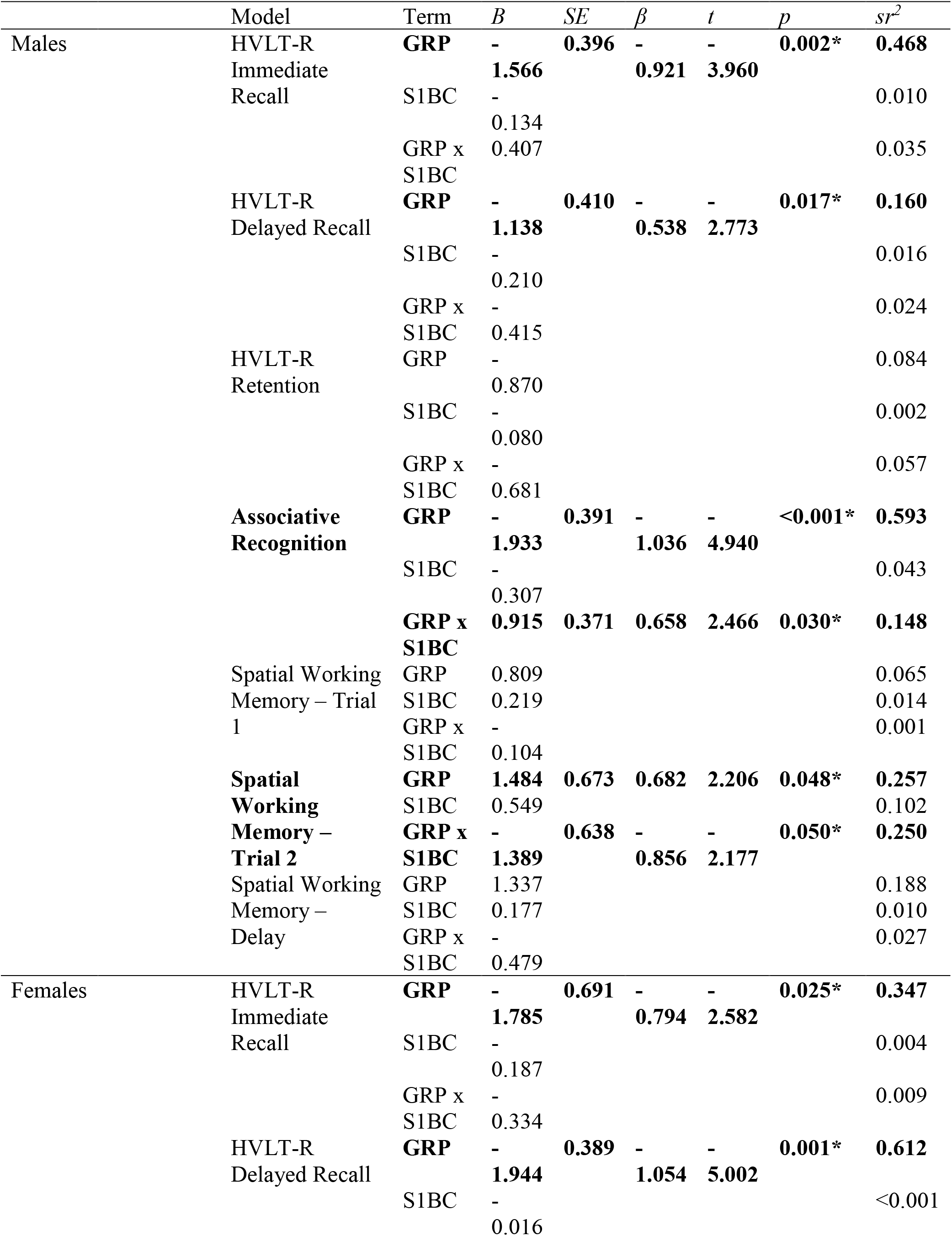
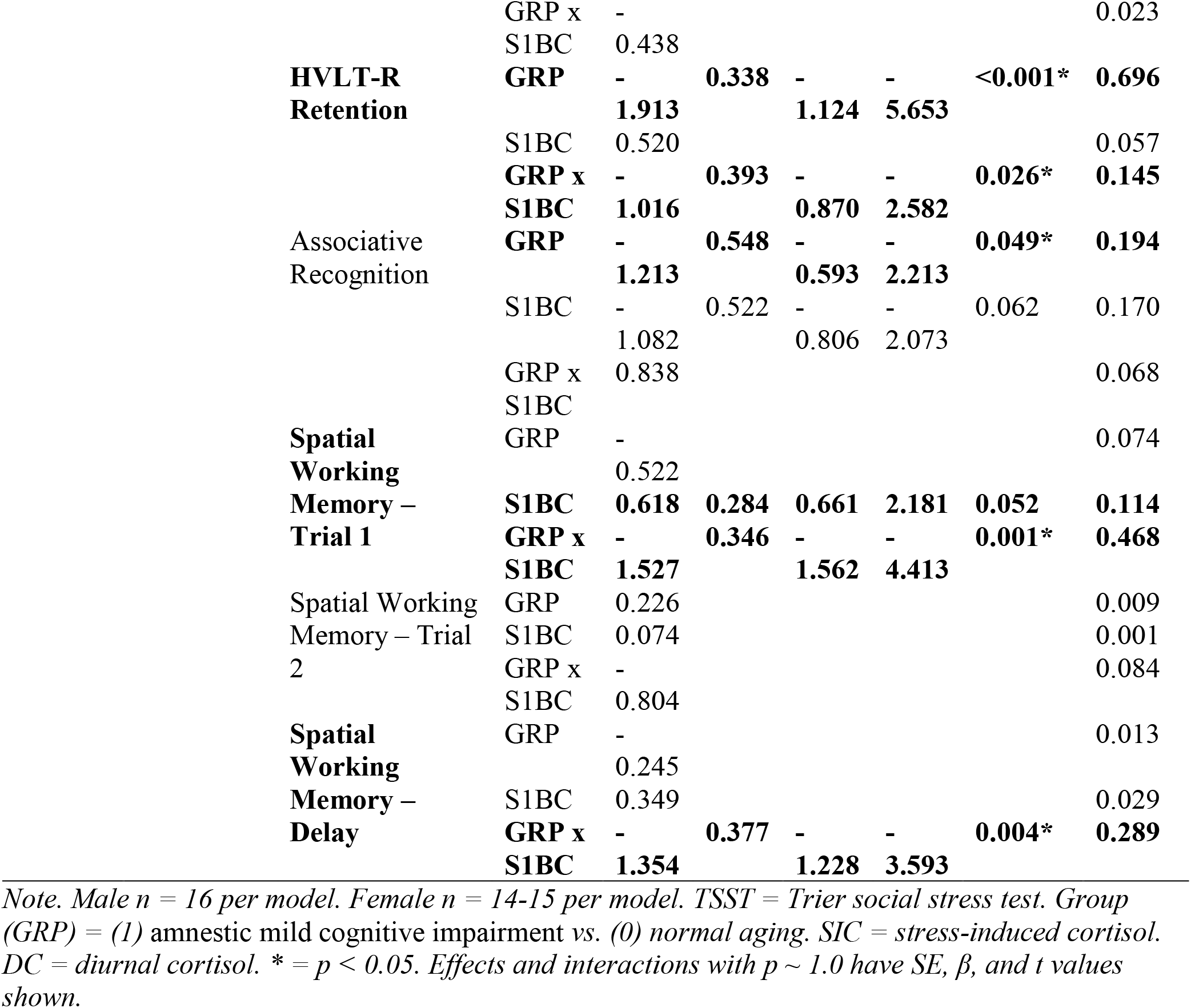
Session 1 cortisol moderation models: main effects and interactions.

### Session 2 cortisol (AUCg) moderates the relationship between aMCI and perceptions of anxiety in response to the TSST speech in males

We tested the hypothesis that the magnitude of Session 2 cortisol (AUCg) moderates the relationship between participant group (aMCI, NA) and episodic (HVLT), associative (face-name), working memory (spatial working memory), or perceptions of anxiety to the TSST speech during Session 2 in males and females. Regressions revealed that high Session 2 cortisol predicted low perceived TSST speech anxiety ratings in aMCI males versus high perceived TSST speech anxiety ratings in NA males (model: *F*_(3,12)_=7.106, *p*=0.005, *R^2^*=0.640, adjusted *R^2^*=0.550; interaction: *b*=-1.118, β=-0.832, t_(15)_=-2.876, *p*=0.014, *sr*^2^=0.248; Figure 7). Similarly in females, the interaction of lower perceived TSST speech anxiety ratings in aMCI females versus NA females with high Session 2 cortisol was close to significant (model: *F*_(3,8)_=8.287, *p*=0.008, *R^2^*=0.757, adjusted *R^2^*=0.665; interaction: *b*=-1.266, β=-0.632, t_(11)_=-2.217, *p*=0.057, *sr*^2^=0.150) (see Figure 7). The interaction also approached significance for HVLT-R delayed recall in males such that high Session 2 cortisol predicted increased Session 2 HVLT-R delayed recall in aMCI males versus no change in NA adult males (model: *F*_(3,12)_ = 22.995, *p* <0.001, *R^2^* = 0.852, adjusted *R^2^*=0.815; interaction: *b* = 0.597, β = 0.368, t_(15)_ = 1.984, *p* = 0.071, *sr*^2^ = 0.049). Additionally, although no interaction was significant, group was a significant predictor for Session 2 HVLT-R immediate recall and delayed recall in females and males and a predictor for Session 2 HVLT-R retention and face-name associative recognition in males (all *p*s < 0.05) (see Table 4).

**Figure 7.**
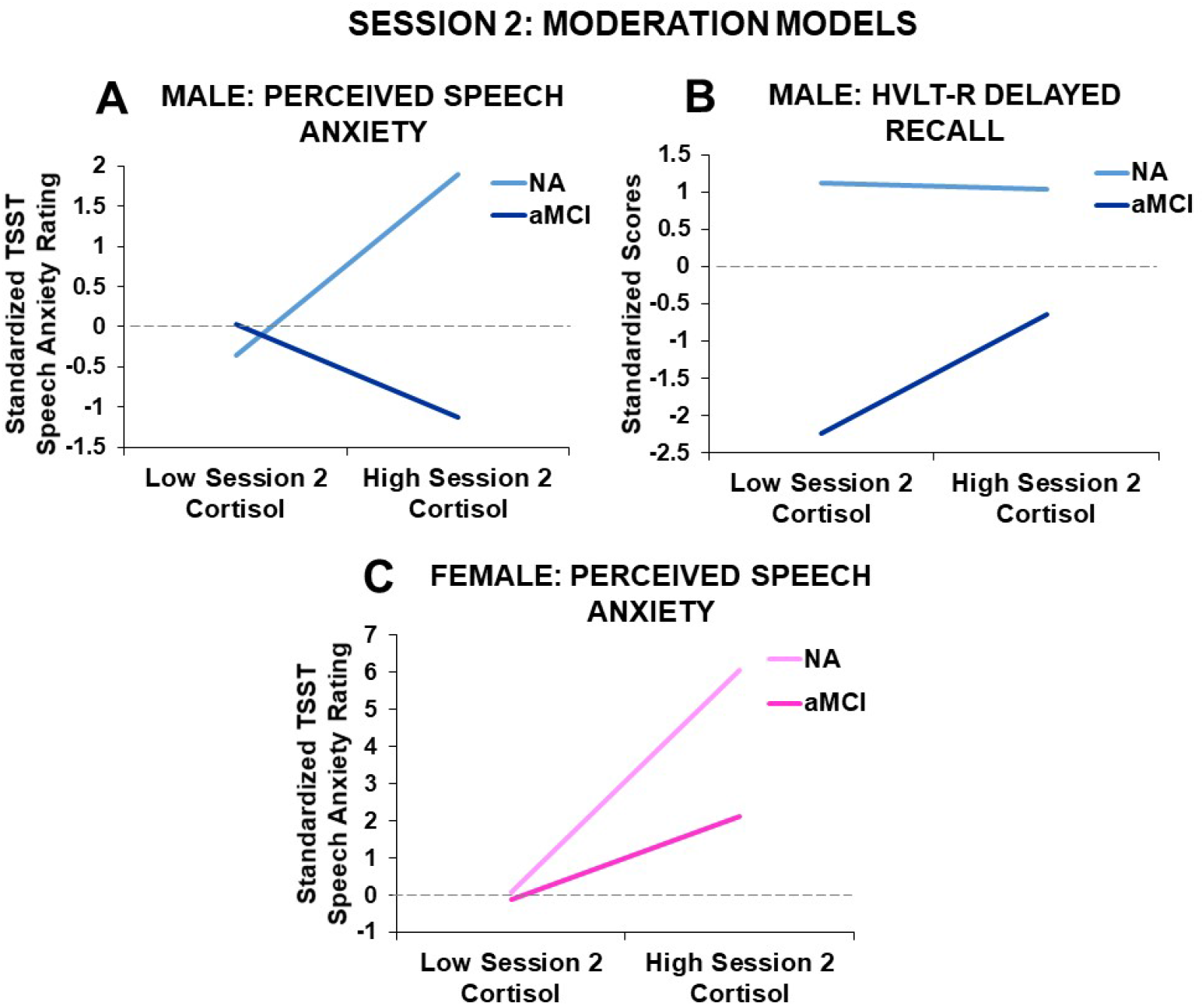
Standardized moderation effects of Session 2 cortisol (AUCg) on the relationship between participant group (normal aging (NA), amnestic mild cognitive impairment (aMCI)) and Session 2 (A) perceived trier social stress test (TSST) speech anxiety ratings and (B) Hopkins Verbal Learning Test-Revised (HVLT-R) delayed recall scores in males (*n* = 7 NA, *n* = 9 aMCI) and Session 2 (C) perceived anxiety rating for the Trier Social Stress Test in females (*n* = 9 NA, *n* = 6 aMCI).

**Table 4.**
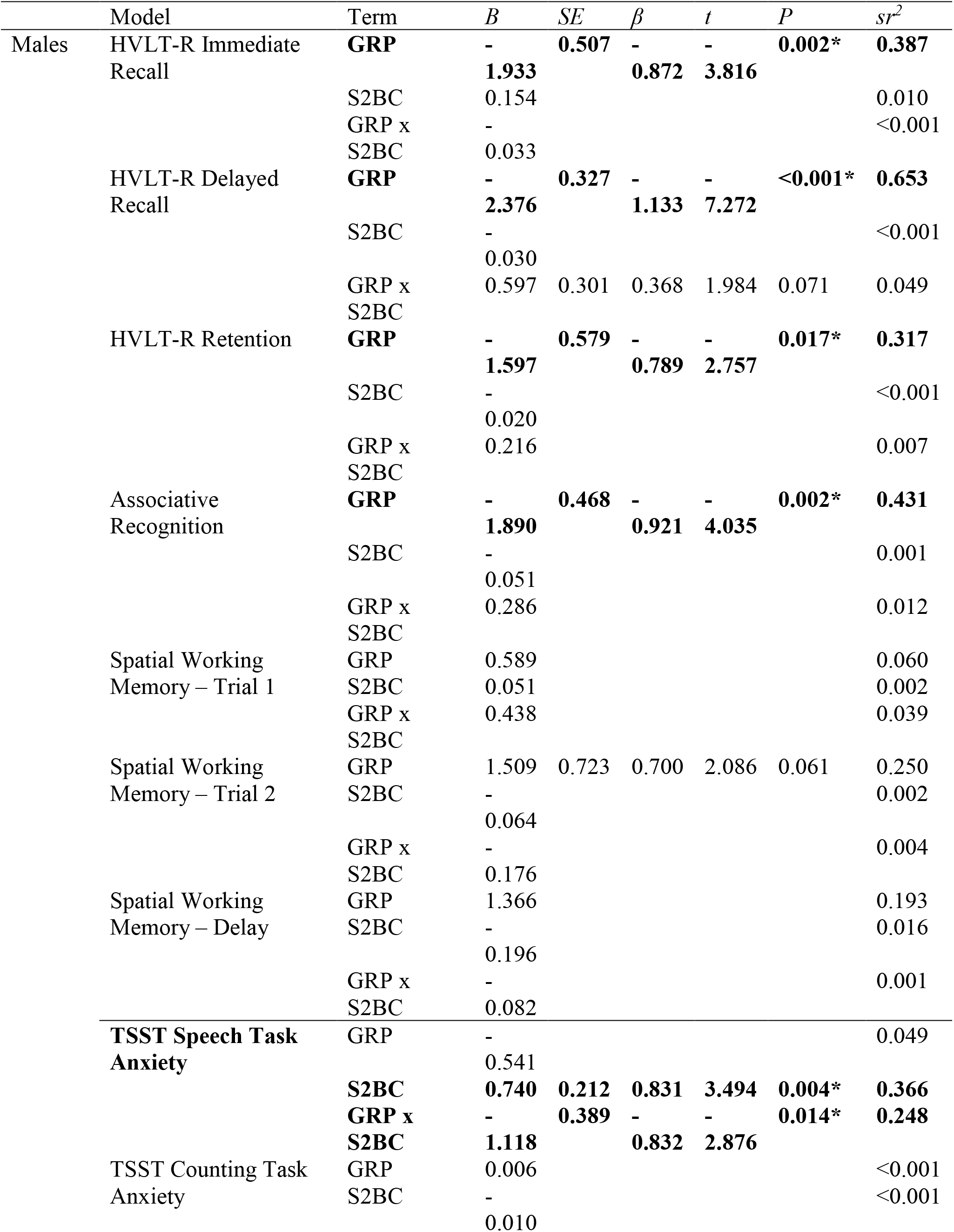
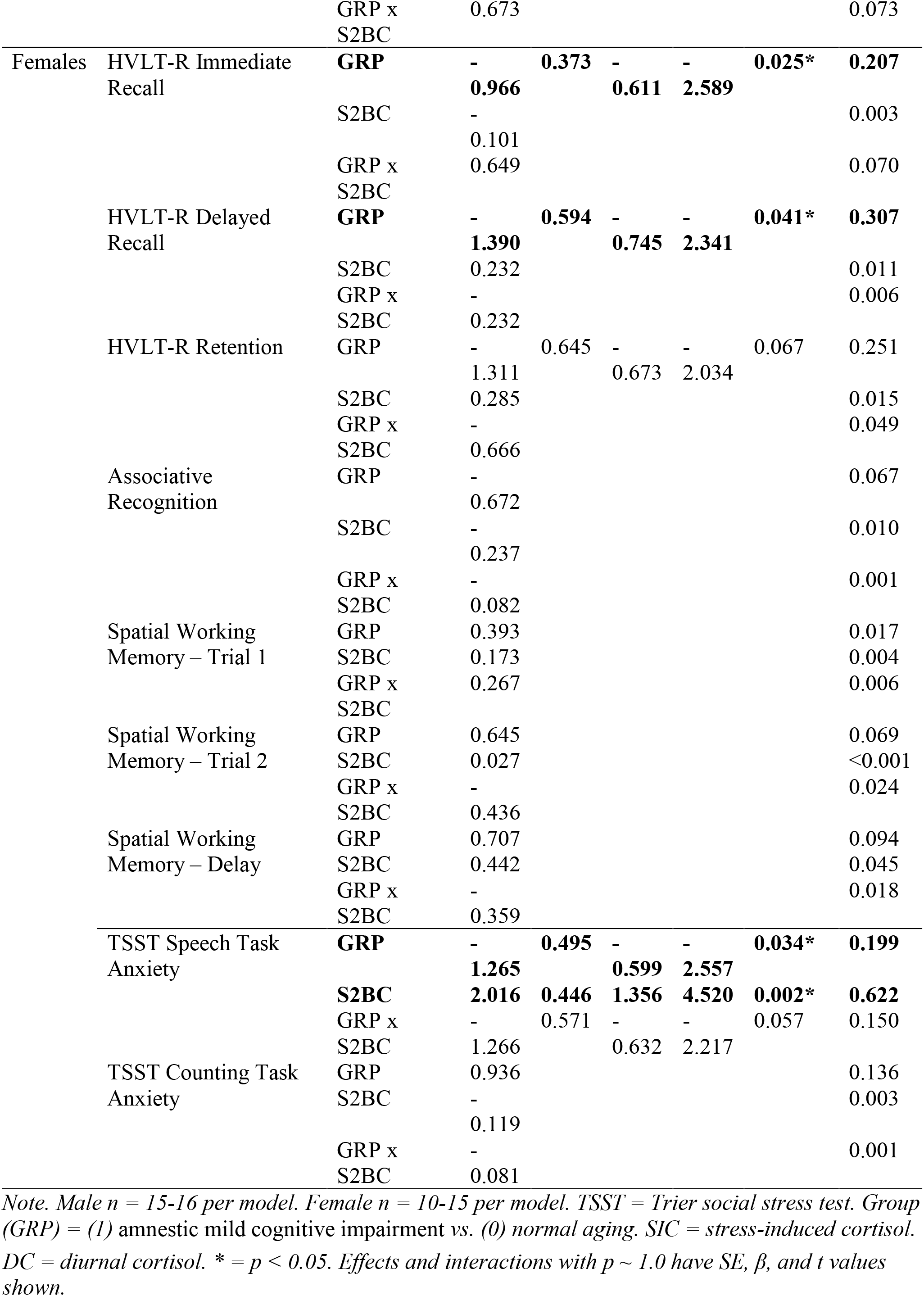
Session 2 cortisol moderation models: main effects and interactions.

## Discussion

Cortisol levels were significantly higher in males with aMCI, an effect that was seen during the test sessions but not in diurnal cortisol, suggesting an effect of the test environment to elicit different cortisol responses among aMCI individuals (consistent with data in NA groups from Sindi et al., 2014). Psychosocial stress, as applied by the TSST, improved immediate verbal recall in NA, but not in participants with aMCI, in fact impairing recall in males with aMCI. Furthermore, our data revealed positive correlations between cortisol levels during Session 1 and spatial working memory in females only with opposing directions based on whether the females had aMCI or NA. Higher cortisol was related to better spatial working memory in aMCI females, but worse spatial working memory in NA females during Session 1. Exploratory moderation models revealed that cortisol moderated perceived anxiety during Session 2 and each of memory domains, dependent on sex and session. Cortisol during Session 1 moderated effects on spatial working memory in both sexes and associative recognition in males, with higher cortisol reducing performance in NA but improving performance in aMCI individuals. Cortisol during Session 1 also moderated effects of episodic memory retention in females such that high cortisol enhanced performance in aMCI but impaired performance in NA females. Stress-induced cortisol in Session 2 was associated with decreased perceived anxiety to the speech in males with aMCI but increased perceived anxiety in NA males, whereas in females cortisol moderated the effect on perceived anxiety positively in both groups but a lower anxiety rating with high cortisol in aMCI females than in NA females. While our sample size was small, our results are suggestive of sex differences in the relationship between cortisol in a testing environment and memory and perceived anxiety that depended on whether the participants were NA or had aMCI. These findings, while exploratory, suggest that sex must be considered when exploring relationships between stress biomarkers and memory.

### Cortisol levels are higher in males with aMCI

Amnestic MCI males had higher salivary cortisol levels as shown in the first samples of both morning sessions conducted in the laboratory. This finding is partially consistent with past studies that found increased morning serum cortisol levels in men but not women with AD (Rasmuson et al., 2011) or in salivary cortisol in individuals with MCI (Venero et al, 2013). In another study, salivary cortisol levels upon awakening were significantly higher in the non-amnestic and multidomain type but not in the aMCI compared to NA (Venero et al., 2013). Even though we found higher morning cortisol in aMCI males in the laboratory setting, cortisol levels did not differ between aMCI males and NA from diurnal samples taken at home, consistent with the Venero et al. (2013) study in which participants also took home samples. Rasmuson and colleagues (2011) found increased morning cortisol in males with AD compared to neurotypical participants they had a low number of participants. Nevertheless, it is compelling that in our findings increased morning levels of salivary cortisol are associated with aMCI in males, at least in the laboratory setting. In support of this, higher cortisol and greater variations in cortisol (and correlations with hippocampal volume) are seen in NA older adults when they were tested in an unfamiliar versus familiar environment (Sindi et al., 2014), suggesting that stressful environments influence correlates of memory. Combined with our findings concerning the influence of stress-induced cortisol on anxiety ratings in males and females with aMCI, our results suggest that further investigation into sex differences in cortisol levels is necessary in individuals with aMCI, and perhaps in the laboratory versus home setting.

What might the possible mechanisms be for the cortisol differences and moderation effects between sexes, and greater impairments in males with TSST on memory? Cortisol is only one output of the HPA axis and other outputs may be important to monitor such as alterations to alpha amylase. In addition, here we have captured acute and diurnal cortisol and it would be important to also examine the influence of allostatic load on these results as well as measures of chronic stress (Yan et al., 2018). In addition, HPA is a regulator of immune and metabolic function, which has been implicated as a driver of or in reaction to AD. Thus, other biomarkers such as cytokines (particularly IL-6) and C-reactive protein (CRP) may be fruitful areas of future testing. Indeed, plasma CRP was decreased in AD and MCI of both sexes and CSF IL-16 and Il-8 differed by sex dependent on APOEe4 genotype (Duarte-Guterman, Inkster, Albert, Barha, Robinson, Galea, 2020). There is a paucity of studies on aging, immune, and other biomarkers that have been implicated in AD with sex as a factor that has elicited calls for action (Mielke et al., 2018) and are findings provide further fodder for these calls.

### Stress-induced increases in cortisol were associated with enhanced episodic memory in NA but not in aMCI

As expected, the TSST induced an increase in cortisol levels, which was associated with enhanced episodic memory (verbal recall) in the HVLT-R in NA but not in aMCI participants. These findings are consistent with a study by Wolf et al. (2002), which found that there were negative correlations between average cortisol and immediate recall of paragraphs in MCI participants but not in NA. Similarly, neurotypical older adults have been found to exhibit a positive correlation between high cortisol and memory performance, whereas aMCI subjects exhibit a negative correlation (de Souza-Talarico et al., 2010). Collectively, the present data and previous findings suggest that cortisol has opposing relationships with memory and recall in MCI versus normal aging that may differ in magnitude and direction by sex. Intriguingly we also found that aMCI males did not mount a stress-induced increase in cortisol with the TSST unlike the stress-induced increase in cortisol in the females with aMCI. These results are intriguing given that de Souza-Talarico et al. (2020) found that greater cortisol reactivity in the TSST was related to cognitive decline characteristic of future MCI after five years and their population was 80% female. This may explain why females with aMCI transition to AD at a greater rate than males with aMCI. The ability of the HPA axis to mount an appropriate stress response may be an important biomarker for AD transition with differences depending on sex.

### Sex may influence the effects stress-induced cortisol on memory in aMCI

In a few of our findings, there were opposing effects of cortisol associations or effects of stress between aMCI and NA dependent on sex. This is intriguing and suggests that sex should be considered in future studies and research on age-related cognitive impairment. This is of particular relevance considering that a number of studies investigating memory and cortisol have had an imbalance of males or females in their test groups (e.g. de Souza-Talarico et al., 2010; de Souza-Talarico et al., 2020; Wolf et al., 2002). Furthermore, inconsistencies in the literature around the effects of acute social stress on memory likely depend on multiple factors including sex and age (for review see Hidalgo et al., 2019; Yan et al., 2018). However, due to a limited sample size in the current work, our sex-based analyses are exploratory and due caution should be paid when generalizing the results.

Our findings of sex differences are congruent with previous studies demonstrating epidemiological, symptomatic, and physiological differences between males and females with MCI. The prevalence of MCI has been found to be greater in males than females, with aMCI as the most common type (Petersen et al., 2010). Furthermore, the incidence of MCI is greater in males than in females (Roberts et al., 2012) and recent studies have uncovered sex-specific risk factors for MCI to AD progression (Kim et al., 2015). Although MCI is more prevalent in males, females exhibit faster deterioration in cognitive and functional measures over time (Lin et al., 2015). Sex differences are also evident in cognitive and neurophysiological decline with AD, as females experience accelerated hippocampal atrophy and cognitive decline with AD (Ardekani et al., 2016; Irvine et al., 2012). These findings, in accord with our data, emphasise the need to account for sex differences in future research in memory and cognition. Intriguingly, decreases in hippocampal volume predict progression to probable AD (and MCI) in women, whereas increases in white matter hyperintensities in men predict progression to MCI (Burke et al., 2019). Optimistically, there is preliminary evidence that cognitive training in those with aMCI is more effective in women than men (Rahe et al., 2015).

### The relationship between cortisol and memory may depend on brain health

Higher cortisol levels should not always be thought of as detrimental to brain function, particularly in the face of acute stress (McEwen, 2019). Indeed, in the present study we found enhanced delayed word-list retention on the HVLT-R in the female NA participants with high cortisol levels in Session 1 under the curve. However, the opposite relationship was found in females with aMCI, as higher cortisol under the curve was associated with reduced retention in the exploratory moderating analyses. These findings are consistent with at least two other studies (de Souza-Talarico et al., 2010; Wolf et al., 2002). De Souza-Talarico et al. (2010) showed a positive relationship between cortisol and delayed recall in NA but a negative relationship in people with MCI. Furthermore, Wolf et al. (2002) showed a negative correlation between average cortisol and immediate story recall in MCI participants but no relationship to average cortisol in NA. It is important to note that opposite patterns were seen in spatial working memory and associative memory, with high Session 1 cortisol associated with improved spatial working memory in aMCI males and females and improved associative recognition in aMCI males but impairments in both associative and spatial working memory in NA participants. During Session 2, high cortisol was associated with enhanded episodic memory (HVLT-R delayed recall) in aMCI males versus NA males, but this interaction had a low effect size. These findings in aMCI participants are in contrast to past findings of impaired cognitive performance in aMCI participants with high cortisol (Popp et al., 2015). However, it should be noted that those studies are based on high diurnal cortisol in aMCI participants and saliva samples were not collected on the same days that cognitive tasks were performed (Popp et al., 2015). Cortisol measures used in our moderation models were based on saliva samples taken in the laboratory throughout the days of Session 1 or Session 2. The impact of cortisol on memory task performance may change when aMCI participants are brought into the laboratory setting. Opposite relationships of cortisol to memory may also reflect brain regions recruited during the task as spatial working memory heavily recruits the prefrontal cortex, episodic memory the medial temporal prefrontal cortex and hippocampus, and associative memory the entorhinal cortex (Courtney et al., 1998; Eichenbaum et al., 2017). Thus, our findings suggest that HPA function may be having opposing effects on memory performance in aMCI groups compared to normal cognitive aging groups.

Acute psychosocial stress improved immediate episodic recall in NA but not in aMCI participants. Although cortisol was not a moderating factor on episodic memory in Session 2, this may be due to a number of factors. Stress activates the HPA axis and it is possible had we measured more timepoints of salivary cortisol we may have seen a moderating effect. In addition, it is important to consider that other biomarkers of HPA activation such as corticotropin releasing hormone (CRH), adrenocorticotrophic hormone (ACTH), or sympathetic activation via the sympathetic-adrenal-medullary system (SAM: alpha amylase, epinephrine, heart rate variability) may have a moderating effect on memory with acute stress. Certainly, it is intriguing that the TSST had an adverse effect on episodic memory in aMCI but a positive effect in NA participants. Indeed, other research has found enhancing effects of the TSST on memory, depending on when TSST was administered relative to memory testing (encoding, retention, recall) that depends on age and sex (Hidalgo et al., 2019). Enhancing effects on episodic memory are see in older (middle-aged) NA women with TSST (Almela et al., 2011). Others have found no effect of cortisol during TSST to moderate working memory in older NA individuals (Pulopulos et al., 2015), consistent in part to our findings that TSST did not influence spatial working memory in the present study. Furthermore, there are well known sex differences in the effects of stress in animal models (Goel et al., 2014). However, the effects of age and stress on learning are not as well studied. In light of these findings, we encourage the research community to make it a priority to examine sex as a factor in analyses of aging and cognition.

### Stress-induced cortisol in Session 2 was associated with greater ratings of anxiety in response to the TSST speech in all groups except aMCI males which were associated with reduced anxiety

Our models revealed that Session 2 cortisol moderated the effect of aMCI on perception of anxiety for the TSST speech in both male and female participants. Greater Session 2 increases in cortisol, the session that involved stress exposure, was associated with greater perceptions of anxiety in NA males and females and to a lesser extent in aMCI females. This finding is similar to previous studies that have found higher anxiety scores in healthy male and female participants exposed to stressors (Ellenbogen et al., 2002). This relationship did not hold for males with aMCI for whom higher levels of Session 2 cortisol were associated with reduced perceived anxiety in our study. Anxiety is found in a high percentage of patients with MCI, subjective cognitive decline, and AD (Banning et al., 2020), but a previous study by Guerdoux-Ninot and Trouillet (2019) found lower perceived stress in response to a Stroop test in male and female AD and aMCI participants compared to NA as the task became more effortful. In the present study, a large percentage of aMCI males did not characterize the TSST speech as anxiety provoking. A blunted perceived anxiety response to stress in aMCI males compared to NA males in the present study could be related to previous findings of increased apathy in male and female AD and aMCI participants compared to nonamnestic-MCI (naMCI) and NA participants (Ellison et al., 2008; Lanctôt et al., 2017). Indeed, past data has indicated that AD and MCI patients differ in the prevalence of symptoms of apathy with a greater prevalence in AD versus MCI groups (Siafarikas et al., 2018). However, these past studies did not analyse their males and females separately and could have missed effects of increased apathy driven by aMCI males. Moreover, these past studies did not examine the effects of cortisol levels on perceived stress or anxiety in aMCI groups. Nevertheless, our findings along with past findings suggest that, whereas increasing stress-induced cortisol is associated with increasing perceptions of anxiety in NA, male aMCI participants are less likely to perceive themselves as being anxious or stressed.

### Limitations

This is an exploratory study given our low sample size and needs to be replicated with a larger population to examine sex-specific effects. Indeed, diurnal cortisol findings in aMCI females were missed because of a lack of samples. Missing diurnal data in aMCI females may reflect sex differences in partner support, whereby partners of females with aMCI may be less predisposed to assisting with remembering to engage in saliva sample collection than partners of males with aMCI might have been. In future, it may be important to measure perceived primary support and partner attitudes to ascertain why samples are missed. In the present study, we had only 2 aMCI females that completed all (or more than 75%) of the diurnal saliva sampling. However, when comparing these two females to the rest of the aMCI group with incomplete diurnal samples, they did not differ in age, education, TSST anxiety ratings, or on episodic or associative memory tasks. In addition to low sample size, and as previously mentioned, taking saliva samples at different timepoints and analysing samples for HPA activation biomarkers other than cortisol (ACTH, CRH, alpha amylase, epinephrine, heart rate variability) may have resulted in different moderating effects of aMCI on memory tasks. Nevertheless, even though cortisol is only one biomarker of stress our results did resemble the findings of other studies that examined the effects of stress on memory function in aMCI versus NA participants. It should be noted that while linear trend lines had the best fit for our data we also attempted polynomial regressions (quadratic, cubic) for associations between cortisol and memory task performances specific to each session in aMCI versus NA and in male versus female participants. In addition to linear associations, some quadratic and cubic associations were also observed but the majority appeared to be driven by outliers. In a future study, a larger sample size in each group would help elucidate whether the association between cortisol and certain memory performance may be non-linear depending on aMCI and/or sex. All in all, a larger sample size would help account for some of the losses in home cortisol sampling and improve the power of our statistics to examine the influence of multiple other factors on our data.

A larger sample size would also permit examination of possible aMCI phenotypes. Other researchers have demonstrated heterogeneity within the aMCI subtype on memory performance measures that may be predictive of progression to AD type dementia (e.g., Sanborn et al., 2017). We chose to focus on the aMCI subtype in order to limit the possible influence of heterogeneous underlying neuropathologies to that of incipient AD. Widening the lens to include other MCI subtypes that are more likely to be of mixed etiology may produce different relationships between cortisol and memory than those demonstrated here. For example, as previously described, Guerdoux-Ninot and Trouillet (2019) found that effort increased perceived stress in NA adults and non-amnestic MCI patients but reduced it in aMCI and AD. Thus, it would be interesting to examine larger sample sizes that include MCI subtypes in a future study in order to further appreciate differences between subtypes and to further explore research demonstrating MCI subtypes may have different phenotypes (Edmonds et al., 2019; Sanborn et al., 2017). Certainly, our data demonstrate it would be important to consider the heterogeneity of aMCI and to determine whether biological sex may be a contributing factor to the heterogeneity.

### Conclusions

The present study found relationships with cortisol (stress-induced and morning session cortisol) and aMCI and moderating effects of cortisol on some domains of memory. As expected, the aMCI participants performed more poorly across the memory measures as compared to NA; however within this we found response patterns influenced by biological sex. We found that males with aMCI had higher cortisol levels in the morning during the test sessions. We also found stress-induced impairment with episodic memory only in males with aMCI. Although our sex-based analyses are exploratory due to the low sample size, sex differences were nonetheless observed. It is critical that future studies explore sex as a biological variable as we have presented evidence herein suggesting that effects at the confluence of aMCI and stress can be obfuscated or otherwise eliminated when males and females are combined instead of being considered separately. For real understanding and advancement to take place in this field, biological sex must be considered and statistically analyzed.

Estimates of the prevalence of MCI in the elderly show high variability, ranging from ~3-42% (Ward et al., 2012), due to differences in study methodology, especially with regards to the sample population (age, ethnicity, education-level, etc.) (Ward et al., 2012). Regardless, there is a health care burden associated with MCI (Ton et al., 2017) as those with MCI are more likely to develop AD (Busse et al., 2003; Lupien et al., 1998) and the health care burden of AD is more severe than that of MCI (Ton et al., 2017). The findings presented here indicate future studies should make examining sex differences (their nature, underlying mechanisms, outcomes, etc.) in aMCI a priority, as well as expand upon the influence of cortisol in aMCI and the interactions between these factors.

## Acknowledgments

We gratefully acknowledge the participants for generously volunteering their time and effort. We thank Triti Namiranian, Angelina Polsinelli, Nicole D’Souza, Diana Smith, Preeyam Parikha, and Tallinn Splinter for assistance with data collection and management and Elizabeth Perez for editorial assistance.

